# A common network of residue-residue contacts underlies peptides’ interactions with MHC class II complex

**DOI:** 10.1101/2025.03.22.644772

**Authors:** Alexander E. Kister, Ilya Kister

**Affiliations:** Department of Neurology, New York University Grossman School of Medicine, New York, NY, 10016, USA

**Keywords:** sequence-structure prediction, antigen, MHC class II, class II-associated invariant chain peptide, peptide binding groove

## Abstract

The formation of a stable peptide-MHC class II complex is a critical step in the adaptive immune response. In this work, we investigate the residue-residue contacts that ’anchor’ the peptide between the alpha and beta chains of MHC II and examine whether such anchoring residue-residue contacts are shared among different peptide-MHC II complexes. We hypothesize that there is a similarity between the map of contacts of antigenic peptides with the alpha and beta chains of MHC II and the map of contacts of the “natural” complex of MHC II with the CLIP - the fragment of the gamma chain. Thus, the CLIP-MHC II complex – specifically, PDB structure 3PDO - was taken as the prototype for peptide-MHC II interaction. To compare the contact maps between the prototype structure and antigenic peptides/MHC II in 14 crystal structures, we developed a ‘unified numbering system’ for residues in peptide-MHC II complexes. Using this unified residue numbering system, we show that approximately half of the CLIP-MHC II residue-residue contacts have analogs in structures that involve different antigenic peptides and different MHC II (HLA-DR, HLA-DQ, and mouse A/B) alpha and beta chains. We present here this common network of contacts that underlies peptide/MHC class II interactions, as well as the structural and physicochemical characteristics of these contacts. Based on these shared characteristics, we propose criteria for the specificity of antigenic peptide loading into MHC II, whereby one can predict whether a particular peptide fragment will bind to MHC II as well as the likely localization of the fragment within the peptide binding groove of MHC II.

## I. Introduction

Loading of digested peptide fragments onto MHC II and the formation of stable peptide-MHC II complexes (p-MHC II) for subsequent recognition by CD4+ T cells is a critical step of the adaptive immune response (Jurewicz and Stern, 2019; Roche and Furuta, 2015). Unlike MHC class I, the MHC class II complex has an open binding site, which allows it to bind peptide fragments of a wider range of lengths and sequences than MHC I. A single allele of MHC II can bind to a very large but restricted repertoire of diverse peptides with high affinity (Barra et al., 2018; Jiang and Boder, 2010). The relationship between MHC II polymorphisms and the repertoire of peptides that can be loaded into a specific MHC II is not fully understood. Although there are many AI-based algorithms to predict whether a given peptide will fit into a specific MHC, such as MixMHC2pred, NetMHCpan, MHCrank, and others (Racle et al., 2019; Nilsson et al. 2023; Lawrence and Ning 2022; Yang et al. 2024), there is still no clear understanding of the principles that underly the mutual compatibility of peptides and MHC II. In our study, we aim to uncover the network of the most important (’anchoring’) residue-residue contacts responsible for the peptide binding within the MHC II canyon. Our analysis is based on the hypothesis that peptides that serve as antigens for MHC II largely retain the same network of contacts as the “natural” CLIP - HLA II complex, in which the alpha and beta chains of HLA class II bind to the HLA class II histocompatibility antigen gamma invariant chain. A fragment of the gamma chain called ‘CLIP’ (CLass II-associated Invariant chain Peptide) binds directly to the binding site of HLA II and is retained within MHC II till antigenic peptide loading (Rocha and Neefjes, 2008). To uncover a shared network of contacts that underlies peptide-MHC II complex formation, we examined peptide contacts with alpha and beta chains of MHC II in 14 different X-ray structures and compared them with the contacts in the prototype structure of the CLIP-HLA II complex. The 14 peptide-MHC II complexes were chosen for their diversity: they include different MHC II molecules and different antigenic peptides with high sequence variability.

To compare residue-residue contact maps, we introduced a ‘unified system of numbering residues’, which allowed us to align CLIP with antigenic peptides and reveal a network of contacts that is in common for all CLIP/Peptide-MHC II complexes. This network of residue-residue contacts, which was found to be similar in all structures, is probably mainly responsible for anchoring the peptide in the MHC II groove and for the specific orientation of the peptide in the MHC II binding site. The characteristics of the residues that are part of the anchoring network, such as residue volumes, hydrophobicity, contact preferences of residue contact pairs, peptide mobility, and the effect of the secondary structure of MHC II alpha and beta chains on the residue contacts, will also be discussed.

### A review of MHC class II structure and function

MHC II molecules (called Human Leukocyte Antigen class II, or HLA II, molecules in humans) are synthesized in the endoplasmic reticulum (ER) and consist of two polypeptide chains known as the alpha (α) and beta (β) chains. These chains bind non-covalently to form an α-β heterodimer, which can be visualized as a ‘canyon’ inside which docks a peptide. One wall of the canyon is mostly formed by two helices of the α chain, and the other wall - by four or five helices of the β chain. N-terminals of both chains form two beta sheets of four beta strands each, which comprise the bottom of the canyon. Inside the canyon lies a peptide-binding groove (PBG), which is open on both sides and can accommodate peptides that are 13–25 residues in length. Both alpha and beta chains are highly polymorphic, especially in regions encoding peptide and T-cell receptor (TCR) binding sites, allowing for diverse peptide presentation. Numerous X-ray crystallographic analyses of MHC II molecules in the Protein Data Bank (PDB) have confirmed that all MHC II structures have similar secondary and supersecondary structures, with some minor exceptions, such as the splitting of the fourth helix into two helices in beta chains. Tertiary structures may vary slightly among different MHC class II alleles, consistent with their primary function of presenting a large variety of processed antigens to CD4+ T cells.

### A review of the invariant gamma chain (Ii) sequence and functions

The Ii chain sequence consists of three parts: the N-terminal cytoplasmic tail, the transmembrane fragment, and the C-terminal extracellular segment, which includes a short (about 15-20 residues) region called ’CLIP’. The process of α, β, and Ii chains complex assembly starts with the formation of a trimer (Ii)^3^, which then associates in the endoplasmic reticulum with three newly synthesized α-β dimers forming a stable heterotrimer (α-β-Ii³). The role of Ii chain in trimeric assembly was recently reviewed (Wang et al., 2024). The Ii chain is also required for targeting MHC II molecules to specific endosomal compartments, such as lysosomes and endosomes. In these compartments, the Ii chain is degraded, leaving only the CLIP peptide bound to PBG until it is replaced by the antigenic peptide (Malcherek et al., 1998).

The CLIP peptide is located between the alpha and beta chains, forming bonds with both chains and thereby maintaining a stable conformation of the peptide-binding groove (PBG). CLIP closes the entrance to the groove like a lid, preventing the binding of intracellular peptides to the MHC II molecule (Roche and Cresswell, 1990; Fortin et al., 2013). In the absence of the Ii chain, an altered α-β dimer conformation is observed, leading to improper peptide presentation and autoimmune reactions (Anderson and Miller, 1992).

CLIP is divided into a central part - the core, and the peripheral regions - Peptide Flanking Regions (PFRs). The CLIP core is usually nine amino acids long and has the most important contacts with the PBG (Sant’Angelo et al., 2002). The PFRs vary in length and composition. They can also interact with MHC II residues, influencing binding affinity, but the number of these contacts is much smaller than in the CLIP core.

## II. Materials and Methods

A total of 15 p-MHC II structures were analyzed, including the prototypical CLIP-MHC II complex. The sequences and PDB codes of these structures are shown in Table 1. Six structures are comprised of various fragments of the gamma chain in complex with human DR and DQ chains, and murine H-2 class II A-B alpha chain and H-2 class II A beta chain. The remaining nine complexes comprise different antigenic peptides in complex with various DR and DQ alpha and beta chains. The crystal structure that serves as the prototype of the p-MHC II complex has PDB code 3PDO: it is a complex of DR alpha and DRB1 beta chains with the CLIP fragment (103-119) from isoform 1 of the HLA class II histocompatibility antigen gamma chain (UniProt P04233-1). The sequences and secondary structures of DR, DQ and murine alpha and beta chains from the analyzed structures are shown in Supplementary Table 1.

**Table 1.**
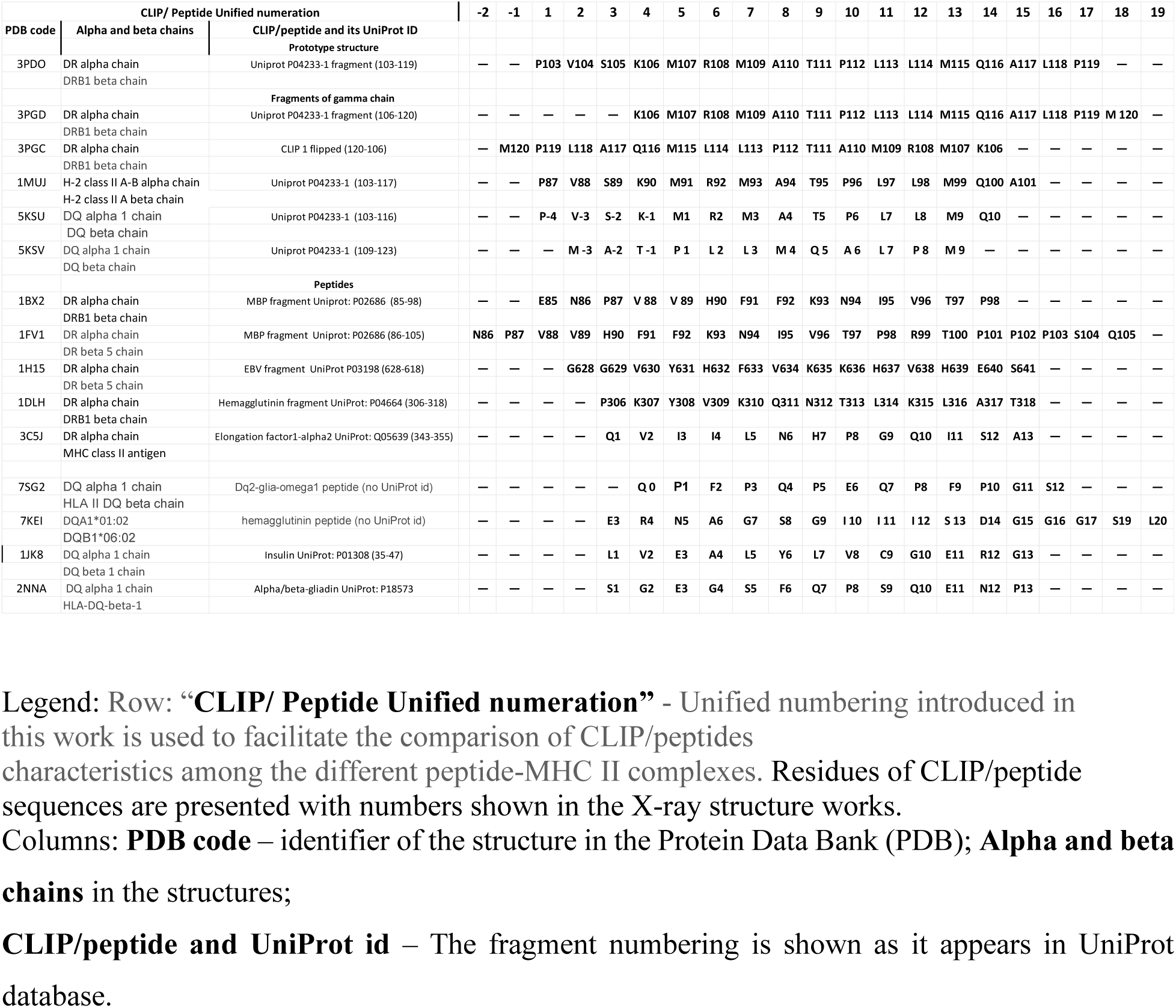
The sequences of variants of CLIP from the gamma chain and antigen peptides.

### Calculation of residue-residue contacts between CLIP or peptide and MHC II alpha and beta chains

To calculate contacts between residues, we used ‘OCA’, a browser-based protein structure/function database that provides contact surface areas (Å²) and hydrogen bond contacts for residues in a molecular complex (Sobolev et al.,1999). The residue-residue contact maps between the CLIP/the antigenic peptides with alpha and beta chains of MHC II are shown in Supplementary Table 2. The area of the contact surface indicates the strength of the interaction: the larger the area, the stronger the interaction.

## III. Results

### Development of the ***‘***unified residue numbering system’ for p-MHC II complexes

The proposed unified system of residue numbering in peptide/MHC II complexes is based on similarities among contact maps of CLIP/MHC II and the antigenic peptides/MHC II. To uncover the similarities we used the following procedure.

1. For each structure, alpha and beta chains were aligned with the alpha and beta chains of the prototype structure (3PDO). Due to the similarity in the number of residues in the secondary structures of the alpha and beta chains in all MHC II complexes, the residues were assigned the same position numbers as the residues in the prototype structure. (as shown in Supplementary Tables 1a and 1b).
2. The residues that comprise the CLIP fragment (103-119) in the prototype structure were assigned sequentially to positions #1-17 in the unified numbering system.
3. The contact maps of all 15 structures were calculated in the unified numbering system for the alpha and beta chains.
4. Comparison of contacts maps of p-MHC II with contact map of the prototype structure revealed that:

a. In CLIP, the residue at position #5 has the largest number of contacts with MHC II: 9 contacts with the DR alpha chain residues and 4 contacts with the DR1 beta chain residues;
b. In every peptide structure, there is one peptide position with the largest number of contacts. The residues at these positions have 7 and 3 contacts with residues at the same-numbered positions in alpha and beta chains, respectively, as does the residue at position #5 in the prototype structure.
5. Given the similarity in contacts between the residue in position #5 of CLIP in the prototype structure and the residues in the highest-contact position of the other peptides, the positions with the largest contacts in all structures were preliminarily numbered as #5. The preceding positions were sequentially numbered #4, 3, 2 … -1, -2, and the succeeding positions were numbered # 6, 7, … 18, 19 so as to form the same continuous numbering sequence. This preliminary numbering of all positions in the peptides must be confirmed by subsequent analysis.
6. The next step was to compare all contacts between residues at the same-numbered positions in the CLIP in the prototype structure and residues in the peptide in the MHC II complexes. We found that there are similar contacts at each consecutive position form #3 to #14 in the CLIP all structures. We call contacts ‘similar’ if residues in CLIP and the peptide with the same position number interact with same-numbered residues in alpha and beta chains, regardless of their chemical nature. For example, Lys in position #4 of the CLIP (prototype structure 3PDO) contacts Ser in position # 53 in the DR alpha chain, while in structure 1JK8, Val in position #4 contacts Arg in position # 53 in the DQ alpha chain (see Suppl. Table 2a). This contact between positions #4 in CLIP/peptide and position #53 in the alpha chain are considered ‘similar contacts’ even though residues in these positions in the two structures are different.
7. To establish a ‘unified numbering system’ for different p-MHC II complexes, we introduce the following requirement: in the unified system, similar contacts must be present in 9 or more consecutive positions in CLIP and peptides. The choice of the minimal number (nine consecutive positions) was made because the CLIP core contains 9 residues (Stumptner and Benaroch, 1997). In our analysis, similar contacts are observed at 12 positions in CLIP/peptide positions. Therefore, the proposed unified numbering system satisfies the above criterion. All Similar Contacts (SC) are presented in Table 2.

**Table 2.**
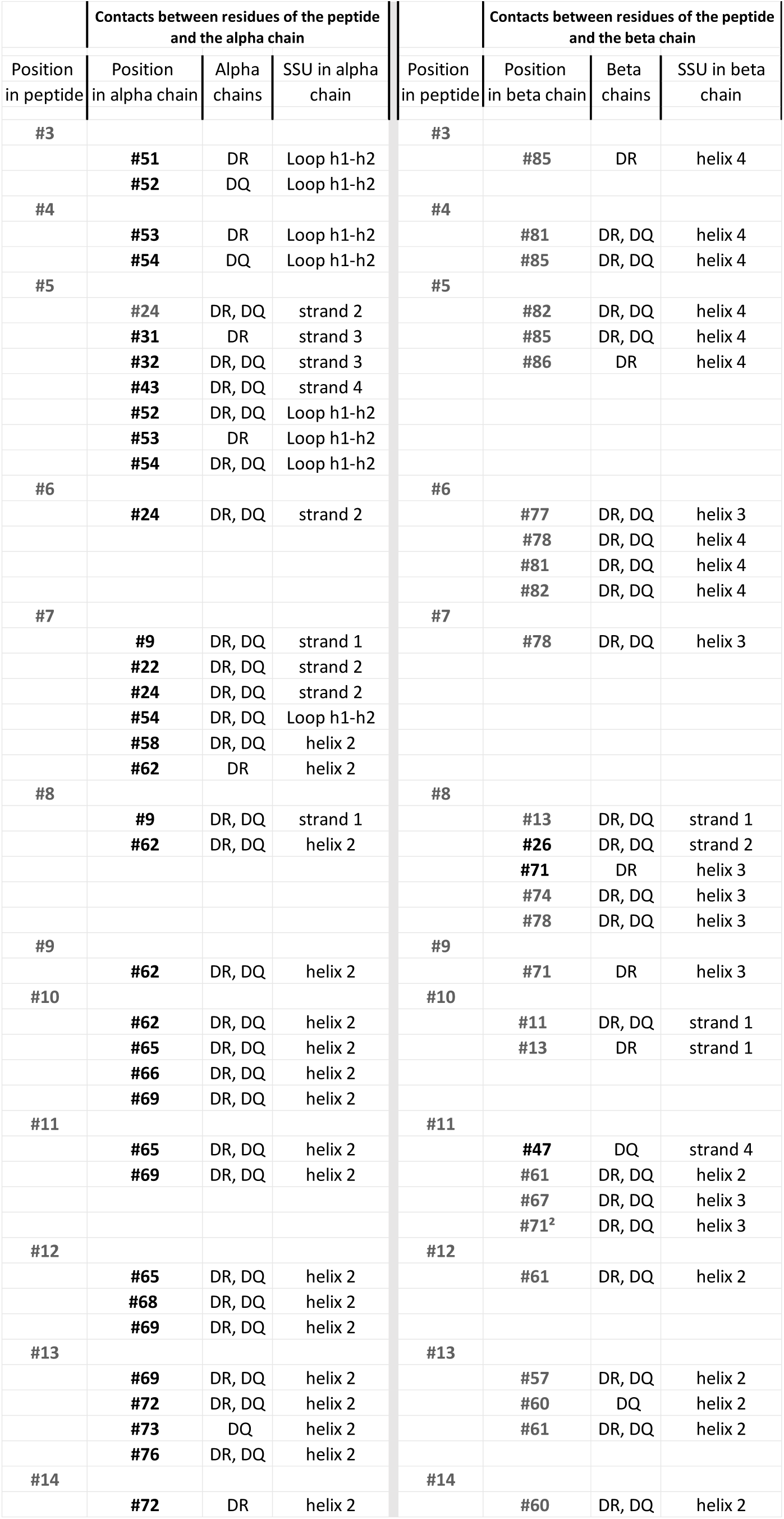
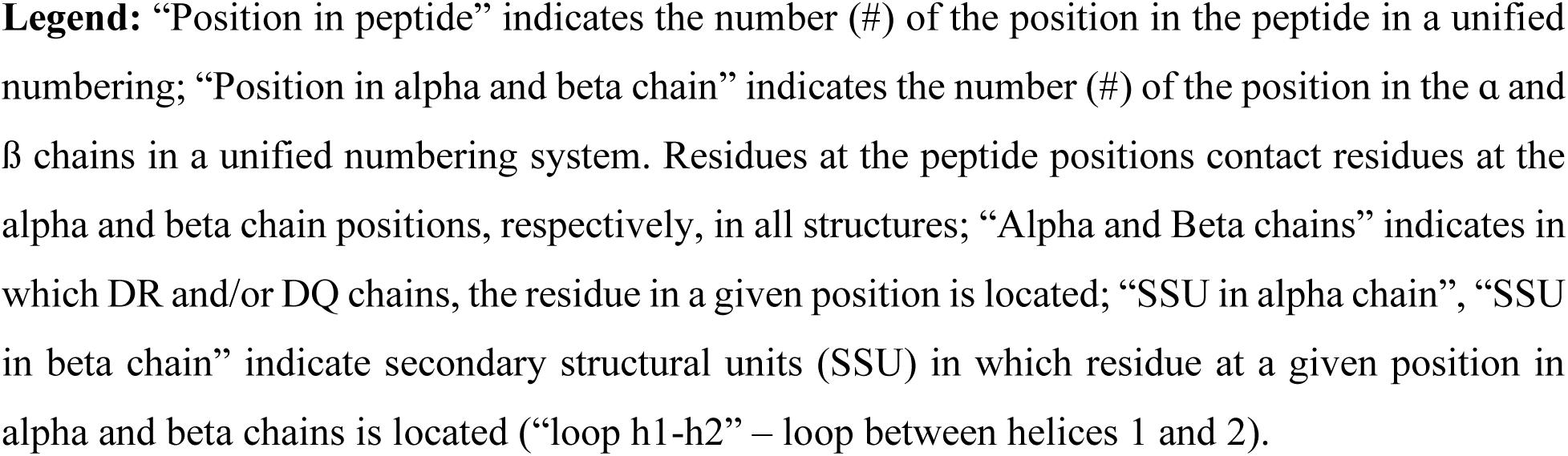
Similar Contacts (SC) between residues of peptide and alpha and beta chains.

| Contacts between residues of the peptide and the alpha chain |  |  |  | Contacts between residues of the peptide and the beta chain |  |  |  |
| --- | --- | --- | --- | --- | --- | --- | --- |
| Position in peptide | Position in alpha chain | Alpha chains | SSU in alpha chain | Position in peptide | Position in beta chain | Beta chains | SSU in beta chain |
| #3 |  |  |  | #3 |  |  |  |
|  | #51 | DR | Loop h1-h2 |  | #85 | DR | helix 4 |
|  | #52 | DQ | Loop h1-h2 |  |  |  |  |
| #4 |  |  |  | #4 |  |  |  |
|  | #53 | DR | Loop h1-h2 |  | #81 | DR, DQ | helix 4 |
|  | #54 | DQ | Loop h1-h2 |  | #85 | DR, DQ | helix 4 |
| #5 |  |  |  | #5 |  |  |  |
|  | #24 | DR, DQ | strand 2 |  | #82 | DR, DQ | helix 4 |
|  | #31 | DR | strand 3 |  | #85 | DR, DQ | helix 4 |
|  | #32 | DR, DQ | strand 3 |  | #86 | DR | helix 4 |
|  | #43 | DR, DQ | strand 4 |  |  |  |  |
|  | #52 | DR, DQ | Loop h1-h2 |  |  |  |  |
|  | #53 | DR | Loop h1-h2 |  |  |  |  |
|  | #54 | DR, DQ | Loop h1-h2 |  |  |  |  |
| #6 |  |  |  | #6 |  |  |  |
|  | #24 | DR, DQ | strand 2 |  | #77 | DR, DQ | helix 3 |
|  |  |  |  |  | #78 | DR, DQ | helix 4 |
|  |  |  |  |  | #81 | DR, DQ | helix 4 |
|  |  |  |  |  | #82 | DR, DQ | helix 4 |
| #7 |  |  |  | #7 |  |  |  |
|  | #9 | DR, DQ | strand 1 |  | #78 | DR, DQ | helix 3 |
|  | #22 | DR, DQ | strand 2 |  |  |  |  |
|  | #24 | DR, DQ | strand 2 |  |  |  |  |
|  | #54 | DR, DQ | Loop h1-h2 |  |  |  |  |
|  | #58 | DR, DQ | helix 2 |  |  |  |  |
|  | #62 | DR | helix 2 |  |  |  |  |
| #8 |  |  |  | #8 |  |  |  |
|  | #9 | DR, DQ | strand 1 |  | #13 | DR, DQ | strand 1 |
|  | #62 | DR, DQ | helix 2 |  | #26 | DR, DQ | strand 2 |
|  |  |  |  |  | #71 | DR | helix 3 |
|  |  |  |  |  | #74 | DR, DQ | helix 3 |
|  |  |  |  |  | #78 | DR, DQ | helix 3 |
| #9 |  |  |  | #9 |  |  |  |
|  | #62 | DR, DQ | helix 2 |  | #71 | DR | helix 3 |
| #10 |  |  |  | #10 |  |  |  |
|  | #62 | DR, DQ | helix 2 |  | #11 | DR, DQ | strand 1 |
|  | #65 | DR, DQ | helix 2 |  | #13 | DR | strand 1 |
|  | #66 | DR, DQ | helix 2 |  |  |  |  |
|  | #69 | DR, DQ | helix 2 |  |  |  |  |
| #11 |  |  |  | #11 |  |  |  |
|  | #65 | DR, DQ | helix 2 |  | #47 | DQ | strand 4 |
|  | #69 | DR, DQ | helix 2 |  | #61 | DR, DQ | helix 2 |
|  |  |  |  |  | #67 | DR, DQ | helix 3 |
|  |  |  |  |  | #71<sup>2</sup> | DR, DQ | helix 3 |
| #12 |  |  |  | #12 |  |  |  |
|  | #65 | DR, DQ | helix 2 |  | #61 | DR, DQ | helix 2 |
|  | #68 | DR, DQ | helix 2 |  |  |  |  |
|  | #69 | DR, DQ | helix 2 |  |  |  |  |
| #13 |  |  |  | #13 |  |  |  |
|  | #69 | DR, DQ | helix 2 |  | #57 | DR, DQ | helix 2 |
|  | #72 | DR, DQ | helix 2 |  | #60 | DQ | helix 2 |
|  | #73 | DQ | helix 2 |  | #61 | DR, DQ | helix 2 |
|  | #76 | DR, DQ | helix 2 |  |  |  |  |
| #14 |  |  |  | #14 |  |  |  |
|  | #72 | DR | helix 2 |  | #60 | DR, DQ | helix 2 |

**Legend:** “Position in peptide” indicates the number (#) of the position in the peptide in a unified numbering; “Position in alpha and beta chain” indicates the number (#) of the position in the $\alpha$ and $\beta$ chains in a unified numbering system. Residues at the peptide positions contact residues at the alpha and beta chain positions, respectively, in all structures; “Alpha and Beta chains” indicates in which DR and/or DQ chains, the residue in a given position is located; “SSU in alpha chain”, “SSU in beta chain” indicate secondary structural units (SSU) in which residue at a given position in alpha and beta chains is located (“loop h1-h2” – loop between helices 1 and 2).

The largest number of SC is observed at CLIP/peptides position #5 – 10 SC. The next highest numbers of SC for the CLIP/peptide positions are in positions #7 and # 8 - 7 SC each, followed by position #10 - 6 SC; positions #6, #11, #13 - 5 SC each; position #12 – 4 SC, position #4 - 3 SC, positions #3, #9, #14 - 2 SC (Table 2).

### Generalized contact map

The unified numbering system of residues of the p-MHC II complex made it possible to align all p-MHC II structures with each other and create a ‘generalized contact map’, which lists the contacts of residues in each position of the CLIP/antigenic peptide with the residues of alpha and beta chains of MHC II. The main conclusions of the analyses of CLIP/peptide - alpha chain contacts in the 15 structures are as follows (Supplementary Table 2a):

> - the residues at peptide positions #1-7 form contacts with the residues of the loop between helix 1 and helix 2 of the alpha chain of MHC II;

> - the residues at peptide positions #2 and #3 form contacts with the residues in helix 1 in the alpha chain in most structures;

> - in all p-MHC II structures, the residue at peptide position #5 forms contacts with the residues in each of the four strands of the alpha chain;

> - the residues at peptide positions #6-8 form contacts with the residues in strands 1 and 2 of the alpha chain;

> - C-terminal residues at peptide positions #7-17 formed contacts with residues of helix 2.

The following is the summary of findings about contacts between residues of the peptide and the residues of the beta chains of MHC II (Supplementary Table 2b):

> - the residues at peptide positions #1-6 always form contacts with the residues in helix 4;

> - the residues at peptide positions #6-11 always form contacts with the residues of helix 3;

> - the residues at peptide positions #11-13 always interact with helix 2;

> - there are no contacts between the peptide residues and helix 1;

> - the residues at peptide position #8 interact with residues in strands 1 and 2;

> - the residues at peptide positions #9 and #11 interact with residues in strands 1, 2 and 4;

> - the residues at peptide positions #10 and 13 interact with strands 1, 2 and 3;

> - in the prototype CLIP/MHC II structure and most other peptide/MHC II structures, there are no residues at positions #15–17.

### Statistical analysis of similar contacts between DR alpha and beta chains and antigenic peptide

The residues at 18 positions of DR alpha chains form 32 SC with antigenic peptide residues (Table 2). Of these 18 positions, 10 positions are hydrophobic. Six of 18 positions (#9, 24, 53, 54, 62 and 72) form 2 contacts each, while the residues at positions #24 (Phe) and #65 (Val) form 3 contacts each, and residues at positions #62 and 69 (Asn at both positions) form 4 SC each (Supplementary Table 3a). The ability of Asn to form multiple types of contacts, such as intermolecular hydrogen bonds and salt bridges, allows it to adapt to different structural contexts, potentially contributing to protein flexibility and dynamics.

Residues at 14 positions of DR1 beta chains form 26 SC, and 6 of these 14 positions are occupied by hydrophobic residues (Supplementary Table 3b). Residues Phe, Trp, His, and Val at positions #13, 61, 81 and 82, respectively, form contacts with residues at two positions each, while residues Tyr and Val at positions #78 and 85, respectively, form contacts with peptide residues at 3 positions each.

### Diversity of peptide sequences

The sequences of the peptides we have studied are very diverse: almost all residues are found within these peptides, with the exception of Trp and Asp. For positions #4 to 13, there are no single or similar residues in the same positions in all structures. Pro residue is the most common residue, found in 12 out of 15 CLIP/peptides at different positions. The largest number of Pro is observed in structure 7SG2: 5 Pro residues (Table 1).

### The hydropathy index of residues of CLIP and peptide sequences

The numbers of hydrophobic and hydrophilic residues also varied greatly. The per-residue Kyte-Doolittle hydrophobicity scale was used to assess the hydrophobicity of peptide sequences (Waibl et al., 2022). Negative values indicate hydrophilic amino acids, and the value of ‘–4.5’ is assigned to the most hydrophilic residues – Arg. Positive values characterize hydrophobic residues with Ile assigned the highest value of ‘+4.5’ as the most hydrophobic residue. The hydropathy indexes for all residues within the peptides are presented in Table 3. In structure 5KSV there are 8 hydrophobic residues and 3 polar residues. The largest number of hydrophilic residues (9) were found in two peptide sequences (PDB codes:1FV1 and 1H15). Calculations of the sum of the hydrophobicity index have shown that peptides can be either predominantly hydrophobic or hydrophilic.

**Table 3.**
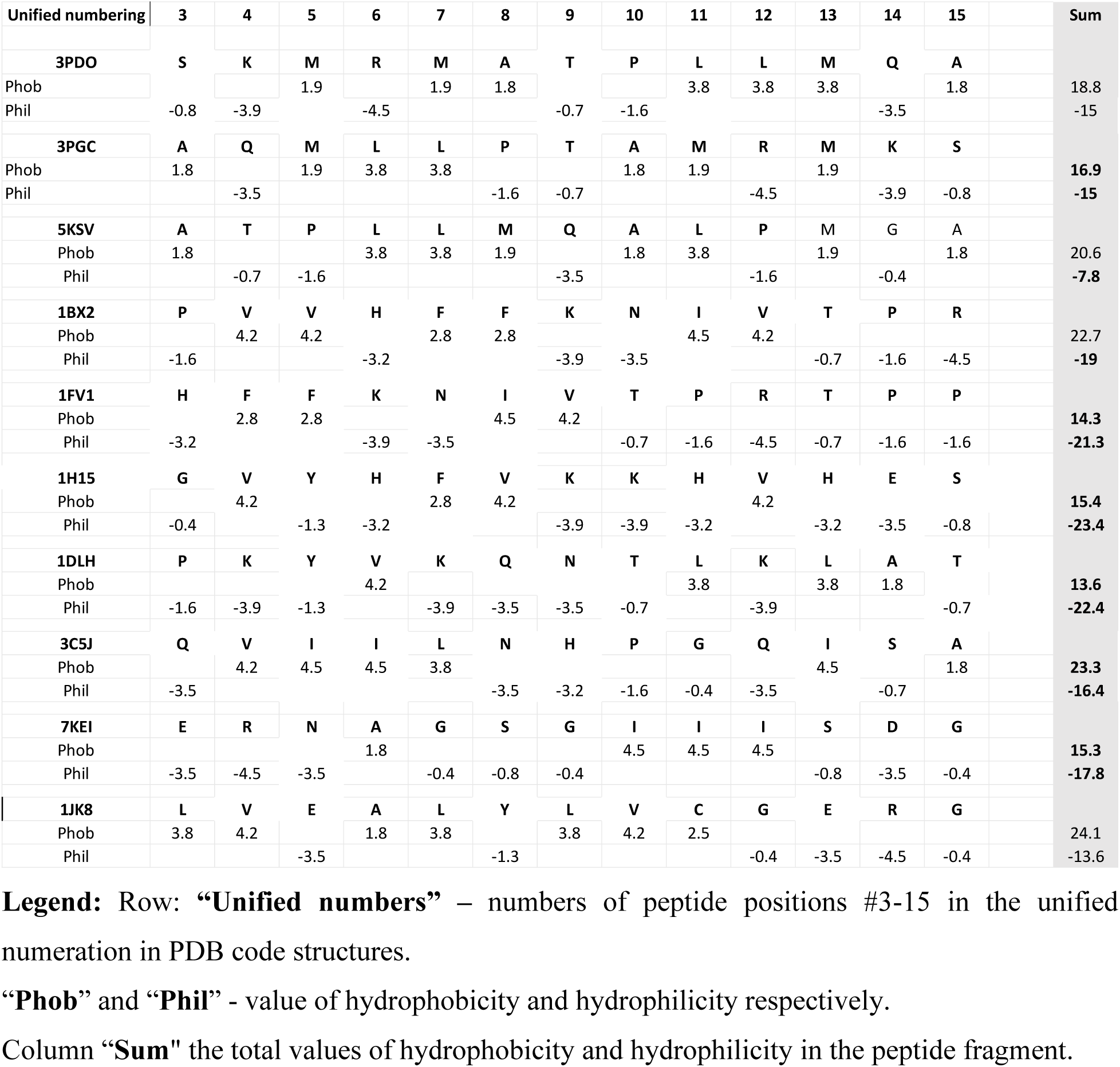
The hydropathy plot of peptide sequences.

Such variability in the hydrophobicity index was also observed for a four-residue fragment within the peptide comprising positions #5-8. This fragment has the most contacts with alpha and beta chains relative to the other parts of the peptide. It can be mostly hydrophobic or hydrophilic, though at least one residue in the four-residue fragment always has the opposite sign from the overall hydrophobicity index of the fragment.

Peptide position #5 can be considered as the only conserved hydrophobic position in the p-MHC II with the DR chains. Although Tyr in position #5 in structures 1H15 and 1DLH has a negative hydrophilic index, it is generally considered a hydrophobic amino acid due to its benzene ring, which repels water.

In all structures except for 5KSV and 3C5J, at least one positively charged residue (Arg or Lys) or at least one negatively charged residue (Asp or Glu) is observed at positions in the peptides. Two structures without charged residues contain 4 polar residues (3C5J) and 3 polar residues (5KSV).

### Volumes of CLIP/peptides’ residues

The volume of amino acid residues is an important factor influencing peptide binding with the alpha and beta chains. The total sums of the volumes of the peptide residues at positions #4 -13 vary from 1392Å³ to 1549Å³ in the structures with the DR alpha and beta chains and from 1165 Å³ to 1402Å³ and in the structures with the DQ chains. The variability of peptide volume values in peptide positions is also very large. For example, the Ala residue (volume 87 Å³) and the Phe residue (volume 190Å³) occupy the same position #4 (Table 4). Residues with small volumes (60-138 Å³) do not occupy positions #4-6, and residues with large volumes (190 and 194 Å³) do not occupy positions #9-13. In contrast to DR chains, the peptide residues with small volumes (60 - 116 Å³) that form contacts with DQ chains were found in peptide positions numbered #4-13.

**Table 4.** Volumes of peptide residues.

| Unified numeration | 4 | 5 | 6 | 7 | 8 | 9 | 10 | 11 | 12 | 13 |  |
| --- | --- | --- | --- | --- | --- | --- | --- | --- | --- | --- | --- |
| Structures | | | | | | | | | | | $\Sigma$ Sum |
| DR structure |  |  |  |  |  |  |  |  |  |  |  |
| **3PDO** | **K** | **M** | **R** | **M** | **A** | **T** | **P** | **L** | **L** | **M** |  |
| Volume Å<sup>3</sup> | 169 | 163 | 173 | 163 | 87 | 116 | 113 | 167 | 167 | 163 | 1481 |
| **3PGC** | **Q** | **M** | **L** | **L** | **P** | **T** | **A** | **M** | **R** | **M** |  |
| Volume Å<sup>3</sup> | 144 | 163 | 167 | 167 | 113 | 116 | 88 | 163 | 173 | 163 | 1457 |
| **1BX2** | **V** | **V** | **H** | **F** | **F** | **K** | **N** | **I** | **V** | **T** |  |
| Volume Å<sup>3</sup> | 140 | 140 | 153 | 190 | 190 | 169 | 114 | 167 | 140 | 116 | 1519 |
| **1FV1** | **F** | **F** | **K** | **N** | **I** | **V** | **T** | **P** | **R** | **T** |  |
| Volume Å<sup>3</sup> | 190 | 190 | 169 | 114 | 167 | 140 | 116 | 113 | 173 | 116 | 1488 |
| **1H15** | **V** | **Y** | **H** | **F** | **V** | **K** | **K** | **H** | **V** | **H** |  |
| Volume Å<sup>3</sup> | 140 | 194 | 153 | 190 | 140 | 169 | 169 | 153 | 140 | 153 | 1601 |
| **1DLH** | **K** | **Y** | **V** | **K** | **Q** | **N** | **T** | **L** | **K** | **L** |  |
| volume (Å<sup>3</sup>) | 169 | 194 | 140 | 169 | 144 | 114 | 116 | 167 | 169 | 167 | 1549 |
| **3C5J** | **V** | **I** | **I** | **L** | **N** | **H** | **P** | **G** | **Q** | **I** |  |
| Volume Å<sup>3</sup> | 140 | 167 | 167 | 167 | 114 | 153 | 113 | 60 | 144 | 167 | 1392 |
| DQ structure |  |  |  |  |  |  |  |  |  |  |  |
| **5KSU** | **K** | **M** | **R** | **M** | **A** | **T** | **P** | **L** | **L** | **M** |  |
| Volume Å<sup>3</sup> | 169 | 163 | 173 | 163 | 88 | 116 | 113 | 167 | 167 | 163 | 1482 |
| **5KSV** | **T** | **P** | **L** | **L** | **M** | **Q** | **A** | **L** | **P** | **M** |  |
| Volume Å<sup>3</sup> | 116 | 113 | 167 | 167 | 163 | 144 | 88 | 167 | 113 | 163 | 1401 |
| **7SG2** | **Q** | **P** | **F** | **P** | **Q** | **P** | **E** | **Q** | **P** | **F** |  |
| Volume Å<sup>3</sup> | 144 | 113 | 190 | 113 | 144 | 113 | 138 | 144 | 113 | 190 | 1402 |
| **7KEI** | **R** | **N** | **A** | **G** | **S** | **G** | **I** | **I** | **I** | **S** |  |
| Volume Å<sup>3</sup> | 173 | 114 | 87 | 60 | 89 | 60 | 167 | 167 | 167 | 89 | 1173 |
| **1JK8** | **V** | **E** | **A** | **L** | **Y** | **L** | **V** | **C** | **G** | **E** |  |
| volume Å<sup>3</sup> | 140 | 138 | 87 | 167 | 194 | 167 | 140 | 109 | 60 | 138 | 1340 |
| **2NNA** | **G** | **E** | **G** | **S** | **F** | **Q** | **P** | **S** | **Q** | **E** |  |
| Volume Å<sup>3</sup> | 60 | 138 | 60 | 89 | 190 | 144 | 113 | 89 | 144 | 138 | 1165 |

Column: $\Sigma$ **Sum** – total amount of volumes of residues at positions #4-13

### Conformational flexibility of peptide

Conformational flexibility largely determines whether a given peptide can fit into the cavity between the alpha and beta chains of MHC II (Ferrante, 2013; Sadegh-Nasseri, 2021). The analysis of conformational flexibility was carried out in accordance with the division of amino acid residues into three groups: highly fluctuating (Pro, Ser, Ala, Gly), moderately fluctuating (Thr, Asn, Gln, Lys, Glu, Arg, Val, and Cys) and weakly fluctuating (Ile, Leu, Met, Phe, Tyr, Trp, and His) residues (Ruvinsky and Vakser 2010). We conventionally assign the residues of the highly fluctuating group the value of ‘2’, the moderately fluctuating group the value of ‘1’ and the weakly fluctuating group - the value of ‘0’. In reality, the assessment of fluctuation requires a more comprehensive evaluation as flexibility is related to many factors, such as hydrophobicity, the position of the residue in the peptide, and the physico-chemical properties of its nearest neighbors (Basu and Bahadur, 2020). For the analysis of peptide interactions with MHC II, when the peptide is “squeezed” into the canyon formed by the α- and β-chains, the choice of adequate indices is even more difficult. However, to assess the tendency and comparative possibility of residues in the peptide, one can use the above scale as a rough approximation (Table 5). We found that CLIP and other peptides have quite a large fluctuation index value in fragments #3-15, with the number of residues from the highly fluctuating group varying from 2 to 7. The residues from the highly and moderately fluctuating groups are located in at least 8 and as many as 12 positions of the peptide sequence. The fluctuating residues in the peptides are distributed unevenly. The largest number of the fluctuating residues are found in the flanks of antigenic peptides. Positions #7, 14 and 15 contain, respectively, 7, 6 and 10 residues from the highly-fluctuation group, while other positions typically contain only 1-3 residues with a high fluctuation index. No residues belonging to the weakly fluctuating group were found at positions #14 and 15.

**Table 5.**
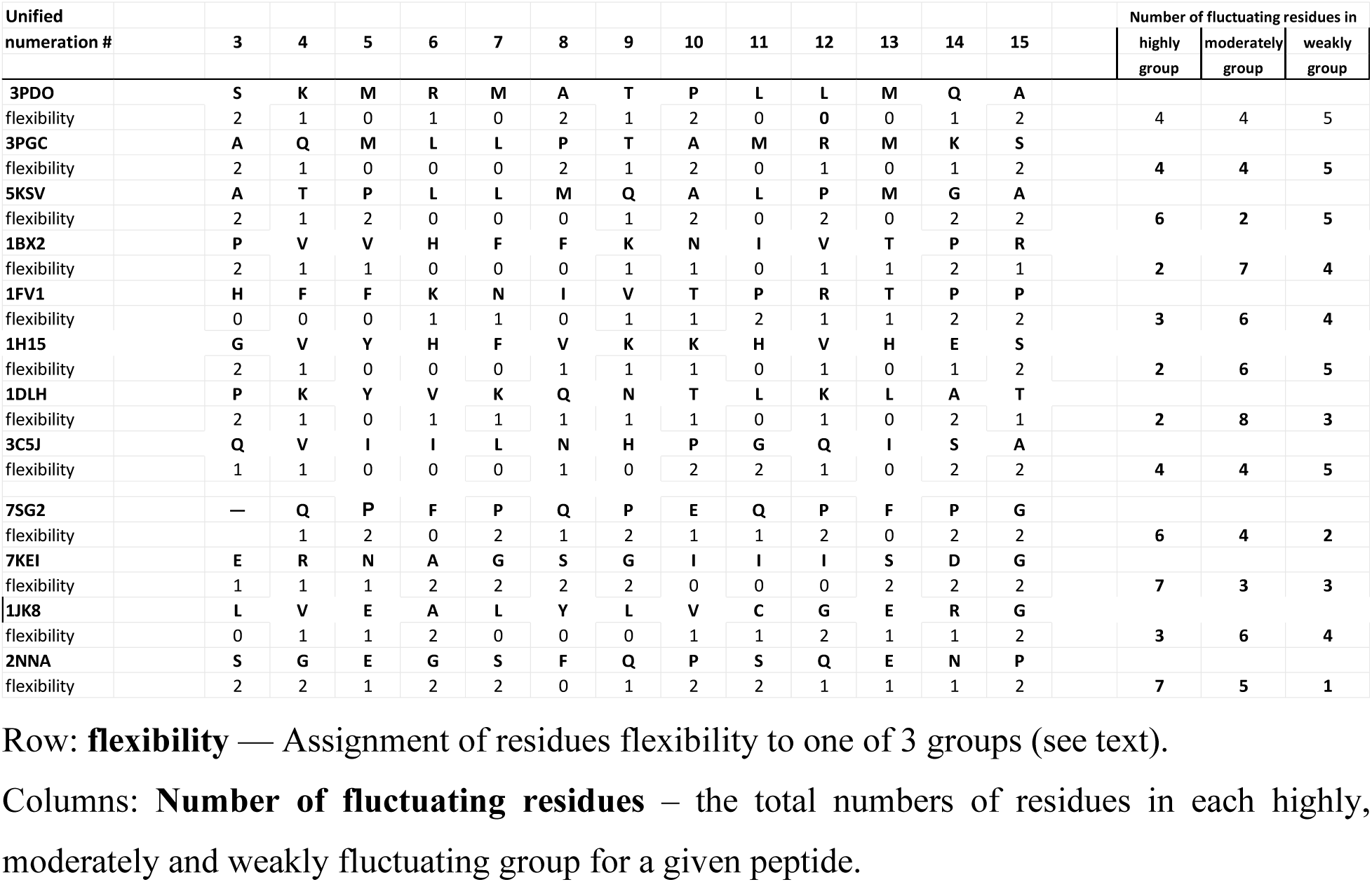
Flexibility of peptide.

### Contact Preference Scores for contacts between peptide and alpha/beta chain residues

The preference score for residue-residue contacts is a quantitative measure used to evaluate the likelihood of interaction between specific pairs of amino acids in proteins. The Contact Preference Scores (CPS) vary from 0 for Cys-Glu contact and 100 for Trp–Trp contact (Nadalin and Carbone, 2017). CPS can be conditionally divided into weak (CPS 0 to 39), strong (CPS 71 to 100) and intermediate (CPS 40-70). The distribution of contacts among these three CPS groups was approximately the same in all structures: a relatively small number of contacts with ‘weak’ and ‘strong’ CPS and a large number of contacts with intermediate CPS (Supplementary Table 3).

To assess the relative contribution of the various peptide positions to peptide’s binding with DR alpha and beta chains, we calculated average CPS values for the similar contacts at peptide positions #3-14 in all the structures with DR chains (Table 6). A comparison of the average values of CPS for similar contacts in each position revealed that:

a. the residues at position #5 of the CLIP/peptide, which have the highest number of contacts, formed contacts with high preference scores (average CPS ≥ 60) in all complexes;
b. the residues at position #7 are characterized by high values of average CPS of ≥60 in all structures except for 1FVI and 1DLH. Similarly, the residues at position #11 are characterized by high values of average CPS of ≥60, except for 1HI5 and 3C5J. Thus, the following observation holds true for all structures: if one of these two positions -either #7 or #11 - has an average CPS of <60, then the other one will have a CPS of ≥60 (Table 6);
c. Positions # 4, 9, and 10 were distinguished by low preference scores (CPS <60) in all structures;
d. For each CLIP and peptide, the sum of the average CPS at positions #3-14 ranged from 652 to 698 (see column ‘O’ in Table 6).

**Table 6.**
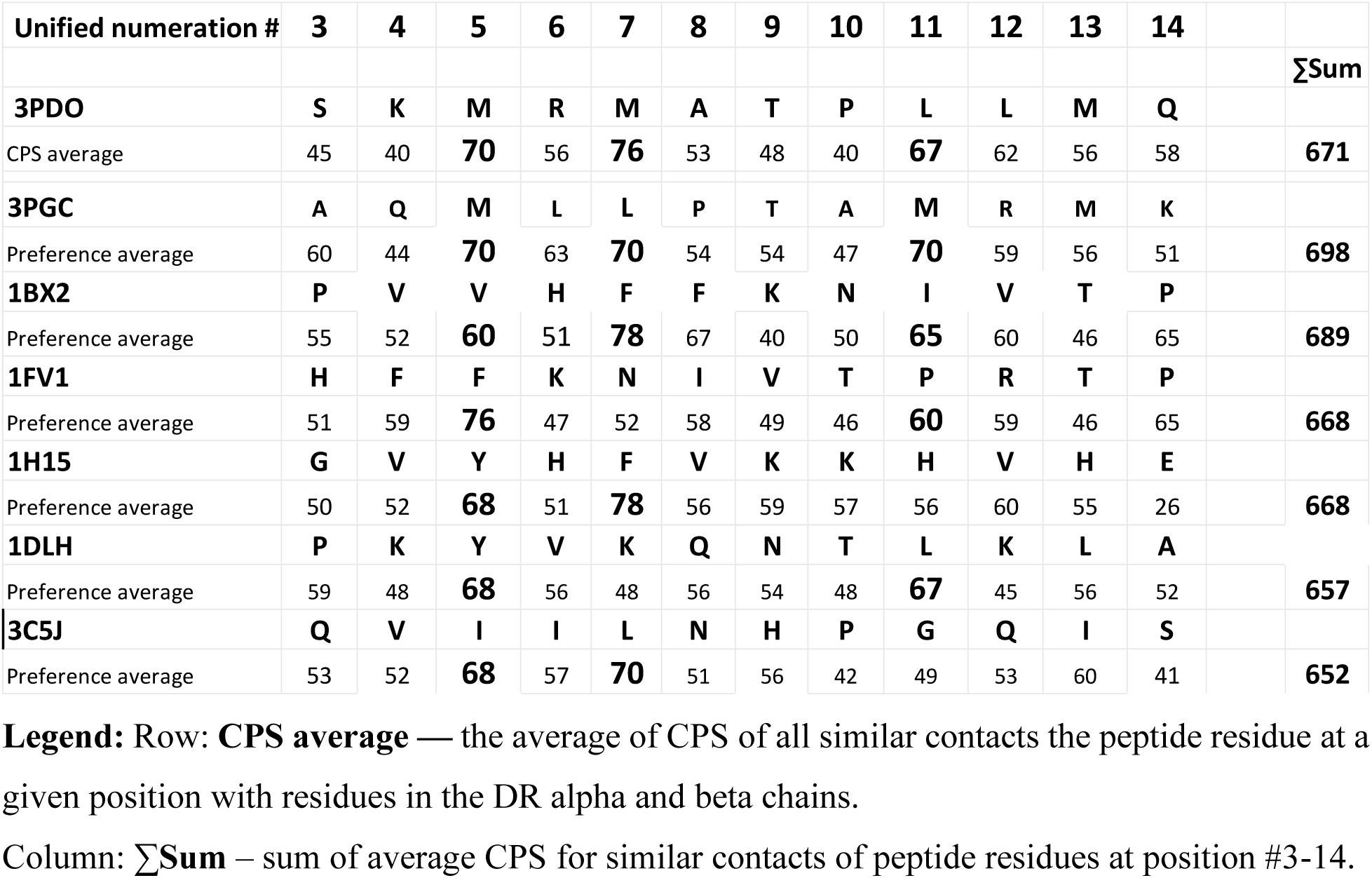
Contact Preference Score (CPS) for contacts between residues of peptides and DR alpha and beta chains.

### Can we use the knowledge of the common contact network to predict which peptides will bind to MHC II?

There is a paucity of reliable structural data on the peptide-MHC II complexes relative to the enormous diversity of peptide antigens that interact with MHC II. However, some predictions about peptide-MHC II interactions can be made on the basis of the common properties of the peptides considered in this paper and the study of contact maps in a generalized numbering residue system.

The question of the interaction of a peptide with the alpha and beta chains can be divided into two interdependent problems. First, can we predict whether a particular peptide has the spatial and sequence characteristics that would allow it to fit into the canyon formed by the alpha and beta chains? Second, if the steric characteristics are satisfactory, can we predict which arrangement of the peptide with the PBG will provide sufficiently stable contacts between the residues of the peptide and the alpha and beta chains? In other words, can we provide an outline of the tertiary peptide-MHC II structure based on sequence characteristics alone without resorting to experimental methods for determining the three-dimensional structures of these complexes?

To answer the first question of the compatibility of peptide and MHC II, we examined the properties of the peptide as a whole: the distribution of hydrophobic and hydrophilic residues, residue volumes, peptide sequence flexibility, and the CPC values. Restricting the ranges of ‘permitted parameters’ for these variables significantly delimits the range of peptides that can interact with MHC II. For example, the analysis of the hydropathy index of residues at positions #3–15 in the CLIP and peptide sequences revealed both hydrophobic residues with positive hydropathy index and hydrophilic residues with negative hydropathy index, but the absolute difference between the sums of positive and negative hydropathy indices did not exceed 5 in any structure (Table 3). Another characteristic of peptides that bind with DR alpha and beta chains is that the residues at position #5 in these structures are always hydrophobic (positive hydropathy index) except for partially hydrophobic residue Tyr.

The constraints on the peptide total volume size follow from the calculated sums of residue volumes at positions #4-13 in the peptide sequence shown in Table 4. The lowest ‘allowed’ residue volumes at position #5 were 140 Å³ in the structures with DR chains. Other bulk characteristics of residues at peptide positions #4-13 can be used to select which antigen is suitable for MHC II presentation.

The flexibility criterion of the peptide can also be used to select an antigenic peptide that will likely interact with MHC II. From the data presented in Table 5, it follows that the minimum number of highly fluctuating residues per peptide is 2. When taken together with the moderately fluctuating residues, the total proportion of residues that contribute to the plasticity of flexible regions is more than 60% of all residues in these peptides.

If most of the peptide compatibility criteria for binding to MHC II are satisfied for a putative peptide antigen, we can tentatively assume that the formation of a p-MHC II complex is possible and consider the second question of localization of the antigenic peptide to the PBGs. The localization of the peptide within MHC II depends on whether it can form a network of stable contacts with MHC II, which is similar to the contacts observed in the 15 structures we have analyzed. Contact preference scores can be used to determine the most likely positions of residues in the unified numeration system for peptide/MHC class II complex. Multiple variants for peptide localization are compared, and the optimal fit is selected as the likely location of the peptide.

The procedure of searching for the optimal localization of the peptide within MHC II is illustrated using the example of structure 3PGS. The sequence of the peptide is presented in Table 7. The procedure for variant #1 is as follows: the residues of the peptide are assigned putative positions in the unified numeration such that the first residue of the peptide (M1) is assigned to the unified position #1, the second residue – to position #2, and so on (see ‘variant #1’ in Table 7). Since all similar contacts for residues at each peptide position with the alpha and beta chain residues have been determined previously (Table 2), we can assign CPS values to all hypothetical similar contacts for residues at positions #3-14 of variant #1. We then calculate the average CPS for each position in variant #1 and compare them to the ‘allowed’ average CPS for the respective positions. The comparison showed that the polar residue Gln is located at position #5, and the average CPS is 55, whereas according to our analysis, only hydrophobic residues should occupy position #5 and the average CPS for residue contacts at this position is ≥60.

**Table 7.** Variants of different localizations of the peptide in MHC II binding site for 3PGC complex.

|  |  |  |  |  |  |  |  |  |  |  |  |  |  |  |  |  |  |  |  |
| --- | --- | --- | --- | --- | --- | --- | --- | --- | --- | --- | --- | --- | --- | --- | --- | --- | --- | --- | --- |
| 3PGS peptide |  | **M1** | **P2** | **L3** | **A4** | **Q5** | **M6** | **L7** | **L8** | **P9** | **T10** | **A11** | **M12** | **R13** | **M14** | **K15** |  |  |  |
| Unified numeration # |  | **-1** | **1** | **2** | **3** | **4** | **5** | **6** | **7** | **8** | **9** | **10** | **11** | **12** | **13** | **14** |  |  |  |
| variant 1 |  |  | **M1** | **P2** | **L3** | **A4** | **Q5** | **M6** | **L7** | **L8** | **P9** | **T10** | **A11** | **M12** | **R13** | **M14** | **K15** |  |  |
| Preference average |  |  |  |  | **79** | 51 | 55 | 68 | **70** | 66 | 52 | 48 | 48 | 62 | 62 | 62 |  |  |  |
| variant 2 |  |  |  | **M1** | **P2** | **L3** | **A4** | **Q5** | **M6** | **L7** | **L8** | **P9** | **T10** | **A11** | **M12** | **R13** | **M14** | **K15** |  |
| Preference average |  |  |  |  | 55 | 54 | 53 | 56 | **76** | 66 | 49 | 42 | 51 | 51 | 56 | 59 |  |  |  |
| variant 3 |  |  |  |  | **M1** | **P2** | **L3** | **A4** | **Q5** | **M6** | **L7** | **L8** | **P9** | **T10** | **A11** | **M12** | **R13** | **M14** | **K15** |
| Preference average |  |  |  |  | **78** | 45 | 68 | 50 | 59 | **70** | 49 | 50 | 55 | 50 | 51 | 62 |  |  |  |
| variant 4 |  | **M1** | **P2** | **L3** | **A4** | **Q5** | **M6** | **L7** | **L8** | **P9** | **T10** | **A11** | **M12** | **R13** | **M14** | **K15** |  |  |  |
| Preference average |  |  |  |  | 60 | 44 | **70** | 63 | **70** | 54 | 54 | 47 | **70** | 59 | 56 | 58 |  |  |  |

**Legend:** Row 1: **3PGC peptide** — Numeration of peptide residues from 3PGC structure

Differences in average CPS values were also found for residues in other positions, e.g., from our antigenic peptide analysis, it followed that the average CPS for residue contacts in position #3 should be < 60, but in variant #1, the average CPS is 79, the highest value for this variant. Another discrepancy with the results of our analysis of peptides in MHC II structures follows from a comparison of the sum of the average CPS contacts of residues at positions #3–14: for variant #1, the total average CPS is 723, which is considerably higher than the total average CPS in the analyzed structures that (between 652 and 698). Thus, the peptide’s arrangement relative to the MHC II binding site in variant #1 doesn’t satisfy the CPS criteria outlined above.

For variant #2, the procedure is repeated, except that the first residue of the peptide now corresponds to position # ‘2’ in the unified numeration scheme. For variant #3, the first residue is shifted one more to position #3, and in variant #4, the first residue is assigned to position #-1 in the unified scheme. These four different variants of the peptide for 3PGC structure are presented in Table 7. A comparison of the average CPS values and other characteristics for the different positions in these variants revealed that only variant #4 yields the average CPS values for the key positions that are compatible with the CPS ranges in the respective positions we have observed in the structures studied in this work. Thus, variant #4 is likely to represent the correct localization of the peptide within MHC II. This indeed is the case: variant #4 corresponds to peptide localization in 3PGS crystal structure.

## IV. Discussion

The criteria that allow for the interaction of peptides and MHC II must, on the one hand, allow for extreme diversity of peptide fragments that fit into MHC II, while on the other hand, also restrict the range of peptides that bind MHC II. The main objective of our study was to identify characteristic of peptide interactions with alpha and beta chains, which are shared by all p-MHC complexes. Our hypothesis is that CLIP contacts with MHC II should be similar to many (possibly most) contacts between peptide and MHC II complexes. Therefore, the CLIP/MHC II complex was chosen as the prototype of the peptide-MHC II complex and the basis for a unified residue numbering system. Our hypothesis was confirmed by the analysis of 14 diverse peptides-MHC II complexes. Three of the complexes we studied involved the same sequence of CLIP as the prototype structure but with different MHC II; two others involved the same MHC II but different ‘versions’ of the CLIP (e.g., shorter CLIP fragment and ‘flipped CLIP’). Peptides in the other nine structures differed widely in their primary sequences from the CLIP sequence. The analysis of the small set of highly variable peptide/MHCII complexes allows one to draw some generalizations about residue-residue contacts in the p-MHC II complex. For example, the lack of hydrophobicity in a position is compensated by the value of contact preference. The positive charge of the Lys residue and the hydrophobic Phe residue at peptide position #4 in the 3PDO and 1FV1 structures, respectively, have similar contacts with Ser residues #53 in the alpha chains with the same contact preference value.

Prior studies have identified shallow dimples, called “pockets,” on an X-ray crystallographic study of the binding sites of HLA-DR1 class II (Brown et al. 1993). The side chains of peptide residues were found to fit in these pockets. However, it is difficult to predict whether a particular MHCII allele will have specific pockets available for binding. Point mutations of residues in alpha and beta chains can affect pocket size and the ability to interact with peptides. In contrast, our work suggests that knowledge of the secondary structure and the ability to represent structures in the unified system of residue numbering may be sufficient to predict whether a specific peptide fragment will form a complex with a specific MHC II. The question of ‘Can the given peptide form a complex with MHC II?’ is reformulated as ‘Can this peptide form similar residue-residue contact with residues of MHC II as a contact map in the unified system of numbering in Table 7’? While it is likely that the small number of input data limits the accuracy of our predictive criteria, it is expected that applying our approach to the increasing number of peptide/MHC II structures will yield an increasingly more precise set of criteria for predicting compatibility of a peptide for MHC II and its localization in a complex. A strength of our work is that the proposed criteria can be readily tested using AI-based computational biology methods.

## Legends to Supplementary Tables

**Supplementary Table 1**. The sequences and secondary structures of alpha chains (Supplementary Table 1a) and beta chains (Supplementary Table 1b)

Residues involved in similar contacts are numbered. They and shown in enlarged size and in bold.

**Supplementary Table 2. The generalized contact map**

Supplementary Table 2a – contacts peptides residues with residues of alpha chains; Supplementary Table 2b – contacts peptides residues with residues of beta chains.

The contacts of CLIP and peptides with alpha and beta chains are divided into three parts: 1. with HLA II DR, 2. with HLA II DQ, and 3 with the murine MHC class II chains. Similar contacts are highlighted in bold.

Columns: **“#pos pept**“: – position numbers of peptide residues;

**# position in alpha or beta chains** – In the column are shown secondary structural unit in the alpha and beta chains and position numbers of residues in these unit, which have contacts with the peptides; In the subsequent columns, for every structure are presented residues that occupy a given position in the peptides and residues in alpha or beta chains which form contact with residues of peptides. In the columns are also shown values of contacts surface area in Å².

The abbreviation means: «no residue» – there is no a residue in peptide at this position in the given structure; “no res. Ch” – means that there is no a residue at this position in the alpha or beta chain in the given structure. “HB’ – stands for Hydrogen bond contact.

**Supplementary Table 3**

**The contact map and contact preference scores (CPS) of DR alpha and beta structures.**

The contacts of CLIP and peptides with residues in the alpha chains (Supplementary Table 3a) and beta chains (Supplementary Table 3b). Similar contacts are highlighted in bold.

In columns **“prefer”** – the contact preference score presents for every contact

